# Single particle tracking with compressive sensing using progressive refinement method on sparse recovery (spt-PRIS)

**DOI:** 10.1101/2022.05.13.491828

**Authors:** Xiyu Yi, Rebika Shrestha, Torin McDonald, De Chen, Harsh Bhatia, Valerio Pascucci, Thomas Turbyville, Peer-Timo Bremer

## Abstract

Single particle tracking (SPT) is an indispensable tool for scientific studies. However, SPT for datasets with a high density of particles is still challenging, especially for the study of particle interactions where the point spread functions (PSFs) are overlapping. In this study, we present spt-PRIS, a new SPT solution where we apply compressive sensing to SPT by integrating the progressive refinement method on sparse recovery (PRIS) into the framework of the state-of-the-art SPT algorithm (uTrack). We systematically characterized and validated spt-PRIS performance using simulations, applied it to the experimental data of membrane-bound KRAS4b proteins in either 2-lipid or 8-lipid membrane supported lipid bilayers (SLB), and compared the results to the conventional method (uTrack). Our results show that spt-PRIS is effective for SPT when the data contains overlapping PSFs and provides unprecedented information about KRAS4b subpopulations. spt-PRIS is helpful for a broad range of scientific studies where precise and fast high-density localization is beneficial. spt-PRIS is also flexible for extensions for multi-species, multi-multi-channel, and multi-dimensional SPT methods with the generalization of PRIS reconstruction schemes.

## 1. Introduction

Single particle tracking (SPT) is a key component in the analysis pipeline in many scientific studies, such as biology and material characterizations [1,2]. For example, SPT is important for the understanding of dynamic protein interactions, where proteins are labeled using fluorophores [1,2] or scattering-based probes [3], and the resulting samples are imaged using optical microscopes to produce thousands of consecutive images, recording the observations of the particle movements over time. The properties of probes (such as locations and brightness) are extracted with single molecule localization methods at each frame and linked across time to produce particle trajectories [4,5]. These trajectories are further analyzed to yield the dynamic information of the labeled particles [2].

In particular, SPT is essential for cancer research, for example, in the study of dynamics of membrane tethered-KRAS4b proteins. Recent studies using SPT have shown heterogeneous and complex diffusion of KRAS4b, which may be associated with dynamic interactions between the protein and lipid microdomains of varying lipid compositions in the plasma memebrane [6,7]. Murakoshi et al. [8] observed transient entrapment of RAS on the cell membrane with increases in the entrapment rate upon stimulation from epidermal growth factor. Lommerse et al. [9] also find similar confinement of RAS within 200 nm. Several studies [10–12] suggest that RAS molecules signal as nanoclusters of 5 to 8 monomers. A recent study by Nan et al. [13] implicates that RAF-RAS-GTP dimerizes in RAF-MAPK activation. The contrasting results about the RAS signaling mechanism as dimers, monomeric GTPase, or clusters consisting of 5 to 8 monomers motivate the need for a detailed understanding of the RAS signaling process and the RAS clustering dynamics in membrane. To address this conundrum, direct tracking of multimer interactions is essential, yet not available. In most of the existing studies, the concentrations of bright molecules are maintained at the single molecule level to assure robustness of downstream SPT analysis. For example, this is done by using a low fraction of labeled KRAS4b proteins [6] or using photoactivation to ensure only a small fraction of the probes are bright at any given moment [14]. Consequently, the fraction of trajectories that directly represent multimer interactions is vanishingly small.

While improving the abundance of multimer-interaction events is not trivial without altering the sample conditions, one could improve the performance of SPT analysis to resolve more trajectories that directly represent multimer interactions. To achieve this goal, three essential methods are required: (1) a *fitting* method to perform robust localization of particles with high local densities that can resolve overlapping point spread functions (PSFs); (2) a *linking* method to produce precise tracking trajectories from the results of the fitting method that may contain rich merging and splitting events; and (3) a *classification* method to distinguish the trajectories enclosing multimer interactions at different scales into different categories.

In this study, we improve SPT by refining the *fitting* method of the SPT analysis pipeline. We apply compressive sensing to the SPT analysis using the recently developed method called the “progressive refinement method on sparse recovery (PRIS)” [15] and demonstrate its ability to provide high-density SPT analysis. We implement the SPT method using PRIS fitting by embedding PRIS results inside the framework of a state-of-the-art SPT package, uTrack [16], and we call the resultant method *spt-PRIS*. Next, we combine spt-PRIS with a recently developed classification method that uses tracking graph (TG) analysis [17] to classify the trajectories into different subpopulations to enable statistical analysis. Sparse recovery (i.e., compressive sensing) is highly effective for precise localization at high densities [15,18]. We utilize PRIS because of its low computational demand, which is suitable for analyzing large-scale datasets for SPT (each contains multiple movies of thousands of frames) while retaining the flexibility for future extensions. uTrack accounts for rich merging and splitting events, and the open-source code package for uTrack allows for fast prototyping and characterization of spt-PRIS by embedding PRIS results inside uTrack. TG analysis [17] is used to mitigate the linking artifacts and to classify trajectories into subpopulations to detailed analysis.

To demonstrate the applicability of the method, we apply spt-PRIS to analyze KRAS4b protein dynamics in supported lipid bilayers with 2-lipids and 8-lipids membrane SLBs, which exhibit low and high clustering effects, respectively [19]. We demonstrate that spt-PRIS results are consistent with prior diffusion-state-based studies and allow access to richer statistics for trajectories showing clustering behavior, providing a significant contribution in achieving SPT for multi-particle interactions. spt-PRIS is broadly applicable to the study of membrane-protein interactions as well as other more general applications of SPT. It can be extended to support 3D particle localization and has the potential for further extensions with multi-channel, multi-species, and multi-dimensional imaging and tracking when integrated with PSF engineering and spectroscopy methods. The ability to directly represent multimer interaction trajectories also motivates further development on trajectory linking and classification for conditions with high-density features.

The rest of the manuscript is organized as follows. Section 2 describes the spt-PRIS methodology, implementation, and the associated TG analysis. Section 3 provides simulation characterizations of spt-PRIS. Section 4 describes spt-PRIS application on experimental data where different KRAS4b clustering dynamics are expected and combined with TG analysis to access subpopulation statistics. Section 5 summarize the method and the results, and discusses directions for future studies.

## 2. Method

### 2.1 spt-PRIS implementation

PRIS is an algorithm for compressive sensing reconstructions (i.e., sparse recovery). Compressive sensing can be used to achieve precise localization of particles when the PSFs of the particles are overlapping due to the benefits of sparsity constraints [15,18]. In PRIS, the solution space for compressive sensing reconstruction is progressively refined to mitigate the computational cost involved with high-resolution reconstructions while retaining the flexibility for various types of generalization options [15].

The spt-PRIS incorporates PRIS in the SPT analysis pipeline by replacing the localization step in the uTrack software [16] with PRIS. In our experiments, PRIS fitting was performed starting on full grids with the original pixel width of 160 nm in 2D to match the experimental pixel size. The fitting was then progressively refined on subsets of pixels with 2-fold refinement for four repetitions to reach a final grid width of 10 nm and, finally, reduced to a list of location coordinates and brightness information using density-based classification [20]. The localized features were further screened to favor SPT datasets with more emphasis on the fitting precision as compared to the recovered fitting density (details are discussed in Section 3.1). The results of PRIS localization were imported into the uTrack software and refined with a least-square fitting routine within the range of 40 nm, slightly above the upper bound of the localization precision for PRIS fitting as shown in Figure 3(a) and discussed in Section 3.1).

The subsequent analyses were performed with the functionality available in uTrack to produce the SPT trajectories. For a fair comparison of localization results, the relevant parameters in spt-PRIS and uTrack analysis were set to be the same. The detected features with location fitting uncertainties more significant than 36 nm (approximately the upper bound of the fitting precision shown in Figure 3(b)) were rejected for both methods to ensure robustness. The SPT trajectories from both spt-PRIS and uTrack on the experimental data were further analyzed with tracking graph analysis (explained in 2.2) to support the relevant statistics analysis on subpopulations.

### 2.2 Tracking graph analysis

Each set of linked SPT trajectories, along with merging and splitting events, form a *tracking graph* (TG), which is a graph that represents correlations between particles across time. In a TG, nodes represent localized features at given time steps, and the edges between nodes represent links between two or more particles at consecutive timesteps.

In our TG analysis, two post-processing operations are involved. First, we identify *loops* in the TG where a node at a given timestep *t* has two outgoing edges to the next timestep that merge back together at time step *t+k*, where *k* characterizes the size of the *loops*. In this study, the *loops* with k<3 are considered to be motion effects and are removed by adopting average particle positions to produce a single continuous track. Given a processed TG, the next step is to compute the *size bound (S*_*b*_) on the number of particles present in any given particle track. We use the method proposed by McDonald et al. [17] to identify a *size bound* for the number of particles in the observed tracks through analysis of the TG. Note that *S*_*b*_ describes the lower bound on the number of particles in the observed tracks due to the presence of unlabeled particles, and a higher bound on the number of bright molecules in each observed feature due to potential bleaching and particle dissociation events. In this way, each feature is associated with a size bound (*S*_*b*_) that classifies the tracks into subpopulations to provide detailed analyses on the properties of the SPT data. The tools are pending approval for open-source release.

### 2.3 Simulations

All simulations used in this study were performed to replicate the common imaging conditions in the study of membrane-bound proteins with the simulation tool used in our previous studies [15,21–25]. The simulations were performed with a grid size of 16 nm in space and 1 ms in time, followed by 10-fold binning to reach a final grid size of 160 nm (to match the pixel size in experiments) and a camera frame rate of 100 Hz. The PSF was simulated with Gibson & Lanni’s 3D PSF model [26] with a full-width half-maximum of 291.1 nm. As shown in Figure 1, the simulations that involve diffusing particles are generated on 10.24 μm wide and 64 nm thick square patches. The Total internal reflection fluorescence (TIRF) field was simulated as an excitation field exhibiting exponential decay from the surface of the coverslip with a penetration depth of 150 nm, and the focal plane was defined to be 16 nm above the coverslip surface to allow for imperfect intensity fluctuation due to particle position changes in the TIRF field. The diffusion trajectories were simulated with Monte Carlo simulation using random walk with time intervals of 1 millisecond. For confined diffusion, the edges of the domains are treated as mirror boundaries. For the full field of view, periodic boundary conditions are used in the lateral directions, and mirror boundaries were used in the axial dimension.

**Fig. 1.**
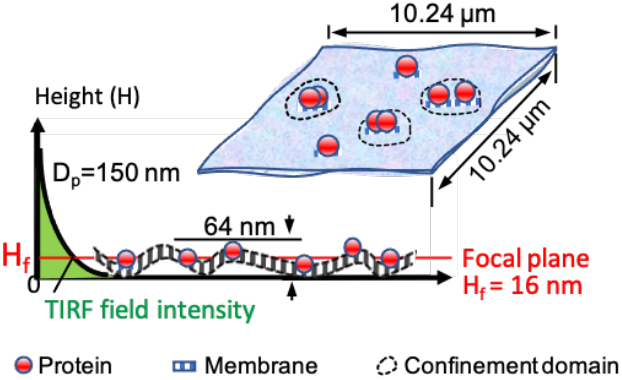
Simulation configuration. The total internal reflection fluorescence (TIRF) field was simulated as an excitation field with exponential decay from the coverslip’s surface with penetration depth (D_p_) of 150 nm. Proteins are distributed in a 10.48 μm wide and 64 nm thick square patch. The focal plane was set at 16 nm above the coverslip. Proteins can have free diffusion in most of the region and confined diffusion in the confinement domains. Merging, splitting, and dissociation events are simulated within the confinement domains.

The imaging processes are simulated with each photon propagated through the imaging system with arrival positions in the detection plane following the probability distribution described by the PSF. Each fluorophore emits a budget of photons weighted by the local TIRF intensity, wherein the brightest condition (at the surface of the coverslip), 500 photons are emitted per millisecond for each fluorophore. Such dependence of the photon budget on the field intensity accounts for the brightness fluctuations due to the changes in the axial positions of the fluorophores. The background photons are simulated as a steady flux of uniformly distributed photons with 20 photons per millisecond per pixel area on average (unless specified otherwise), mimicking the relevant experimental observations. All the noise upon acquisition were simulated as shot noise with Poisson statistics. Note that the simulated protein dynamics do not represent the precise molecular dynamics of KRAS4b protein. Instead, the focus of our simulations is on producing realistic microscopy simulations to represent the imaging imperfections that impedes on the robustness of the localization step in SPT.

### 2.4 Experiments

We prepared two types of supported bilipid layers for single particle tracking of KRAS4b, either 2-lipid or 8-lipid membrane SLBs, and compared the performance of spt-PRIS analysis to the conventional method (uTrack). Supported lipid bilayers (SLB) were prepared as described in the previous study [19,27]. Briefly, small unilamellar vesicles were deposited on a plasma cleaned #1.5, 40 mm borosilicate coverslip and incubated at room temperature for 30 minutes, followed by washing with 20 mM HEPES, 200 mM NaCl buffer. 1 μM of untagged fully processed recombinant KRAS4b proteins mixed with 50 nM Janelia Fluor (JF) 646 [28] labeled KRAS4b S106C was added to the SLB and incubated for an hour at room temperature. The excess unbound proteins were removed by flowing through the imaging buffer (20 mM HEPES, 300 mM NaCl, 5 mM MgCl2 and 5 mM beta-mercaptoethanol (BME)). BME is essential to avoid protein crosslinking via photosensitized oxidation [29]. Detailed protocols can be found in the prior study [27].

Single-particle tracking (SPT) experiments were performed on a Nikon NStorm Ti-81 inverted microscope using 100× oil immersion TIRF objective with numerical aperture of 1.49 N.A. 1.49 (Nikon, Japan). Images were captured with an Andor iXon EMCCD camera (Andor, USA) with 647 continuous wave excitations (Agilent laser module). The samples were briefly photobleached with high laser power to photobleach the fluorophores adhered to the coverslip surface that may skew the diffusion results and immediately imaged at the 20% power setting for 5000 frames for each dataset.

## 3. Validation and characterization of spt-PRIS using simulated data

Seven sets of simulation data were used to validate and characterize the performance of spt-PRIS. The results are compared with those from the conventional method (uTrack) and the ground truth analyzed directly on the input particle trajectories retrieved from the diffusion simulations.

### 3.1 PRIS yields better fitting on clusters

We first simulated clusters of 2 to 8 bright emitters uniformly positioned along the circumference of a circle with a radius ranging from 1.6 μm to zero with intervals of 16 nm, representing a series of increasingly challenging localization tasks for clusters (simulation set 1). Results from PRIS and the localization routine embedded in uTrack are compared. Figure 2 demonstrates the representative results for each cluster size, where PRIS demonstrates faithful localization results in all demonstrated cases, while for the uTrack embedded fitting results, a cluster of emitters is fitted either into a single emitter or into multiple emitters but with larger errors. The complete set of results are shown in visualization 1, which shows both PRIS and the uTrack embedded fitting perform well in the less challenging conditions while PRIS performs better in the more challenging conditions.

**Fig. 2.**
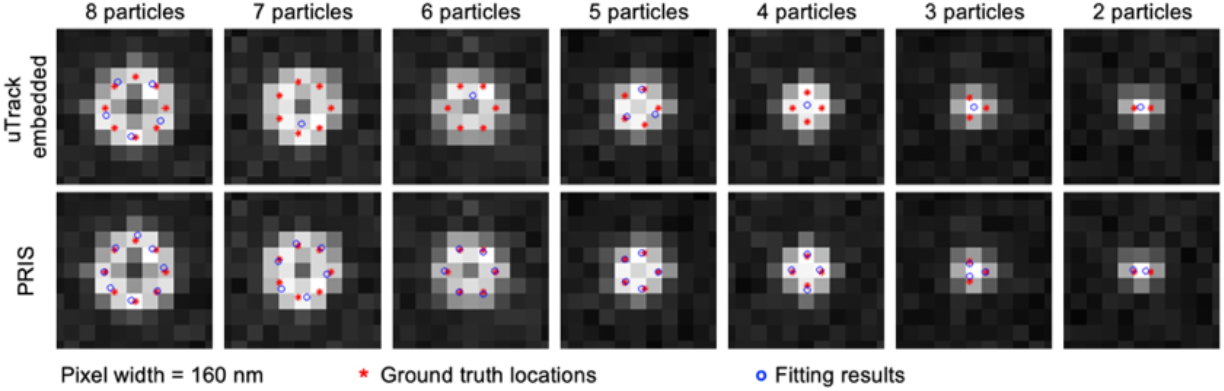
Localization performance on clusters (simulation set 1). Various particles located with different mutual distances are simulated with a background photon count of 40 per pixel area per millisecond. The top row shows the results of localization fitting embedded in uTrack, and the bottom row shows the result from PRIS. Pixel width is 160 nm. The complete set of results are shown in visualization 1.

Next, we characterize the localization performance of PRIS at various particle densities (simulation set 2). The PRIS fitting parameters are configured to suit SPT, where performance on localization precision and false-positive rates are favored over recovered density. We characterized the fitting performance based on three metrics: (1) the localization error, which is the standard deviation of the localization errors; (2) the recovered density, which is the density of the ground truth particles that are identified by the localization method; and (3) the false positive ratio, which is the fraction of localized particles that do not have a ground truth particle located within 200 nm radius of its position. As shown in Figure 3, PRIS demonstrates a better performance than the results from the localization routine embedded in uTrack with lower localization error and higher recovered density. Both methods exhibit a negligible false-positive ratio.

**Fig. 3.**
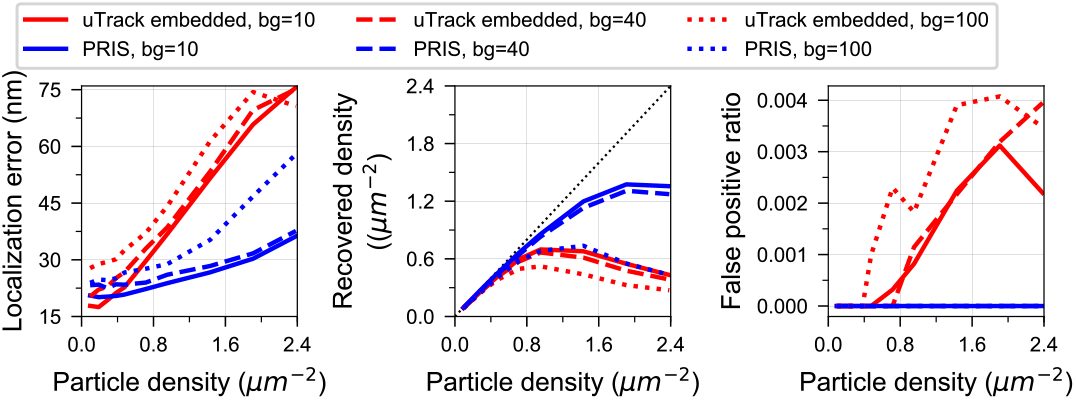
Characterization of localization performance with different particle densities (simulation set 2). Fifty images with emitters randomly distributed in a patch of size 10.24 × 10.24 μm^2^ were analyzed to produce each data point for both PRIS and the localization method embedded in uTrack (uTrack embedded). Three different noise conditions were used for comparison, where the average count of background photons per pixel per millisecond (bg) was set as 10, 40, and 100, respectively as annotated.

### 3.2 PRIS improves linking via improved fitting

Next, we demonstrate that the enhanced localization performance in PRIS further improves the tracking. We simulate four pairs of diffusing particles (simulation set 3), which diffuse at 0.5 μm^2^/s when far apart initially. When they come close, they are locked in at 200 nm with a shared diffusion constant of 0.05 μm^2^/s. As shown in Figure 4, when the two particles are close to each other (Figure 4(a-c)), the fitting routine embedded in uTrack identifies the cluster as a single particle, whereas PRIS recognizes two particles. Additionally, when the subframe motion effect exists for fast diffusion trajectories (Figure 4(d-f)) where the positions of the particle could spread over more than one pixel over the camera exposure time, uTrack returns the average particle position, whereas PRIS fitting results are presented with awareness of such motion effect.

**Fig. 4.**
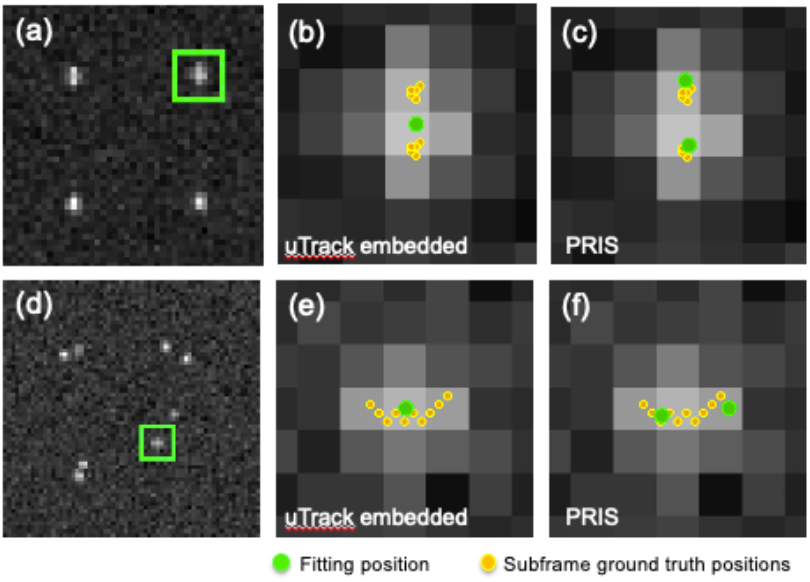
Localization (simulation set 3) comparison. PRIS can identify separate particles and motion effects even at low resolution where uTrack either fails to resolve the particles or fail to recognize the motion effect. Top row: localization with particles with overlapping PSFs. (a) shows a single frame, (b) shows the result from uTrack, and (c) shows the result from PRIS. Bottom row: localization with fast-moving particles with motion effect. (d) shows a single frame, (e) shows the result from uTrack, and (f) shows the result from PRIS. Pixel size is 160 nm. The complete set of results are shown in visualization 2.

Here, we note that the motion effect is not accounted for in the standard linking algorithms (uTrack). Such events would be identified as separate particles and cause the linking algorithm to assign separate trajectories to the corresponding movement, resulting in trajectory artifacts. Although it is conceivable that the motion effect can be identified based on the time duration of the particles and the transient changes in the particle intensities, and an improved linking algorithm could be developed with an awareness of the motion effect, it is beyond the scope of this study. Here, we handle the motion effect using the trajectory cleaning method discussed in Section 2.2.

Figure 5 shows the localization results over the full time-range (250 frames) for simulation set 3, where the close-range interactions of the particles exist from the 100^th^ to 150^th^ frame, as can be seen in the ground truth particle position plot (Figure 5(a)). We note that spt-RPIS (Figure 5(b)) identifies two individual particles in these clusters that match the ground truth closely, whereas uTrack embedded fitting results (Figure 5(c)) identifies the clusters as single particles. Such a condition is critical when one needs to study the diffusion statistics of the subpopulation of clusters with different sizes, and our result suggests that PRIS provides the localization that more faithfully represents the particle trajectories under such conditions.

**Fig. 5.**
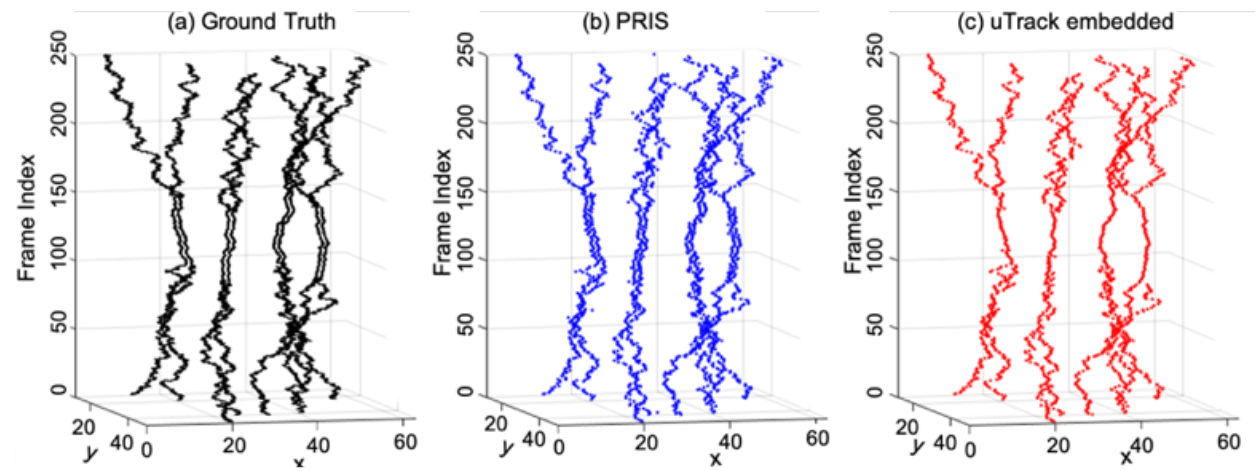
The plot of the fitted location demonstrating the diffusion traces (simulation set 3). (a) shows the input particle positions used to generate the microscopy observations. (b) shows the recovered position using PRIS fitting. (c) shows the recovered position using the fitting routine embedded in uTrack.

We further increased the complexity of the simulation to involve mixtures of individual diffusing particles and pairs of particles with random, close-range interactions (without merging) to confirm the robustness of spt-PRIS (Simulation set 4, Figure 6). The simulated sample contains 30 particles with diffusion constants of 0.01 μm^2^/s, 0.03 μm^2^/s, and 0.09 μm^2^/s (10 particles each), and an additional ten pairs of particles (located 200 nm apart for each pair) with 0.01 μm^2^/s translational diffusion and 0.0152 rad^2^/s rotational diffusion (the root mean square angular displacement per 10 ms is 1 degree). SPT analyses were performed using both spt-PRIS and uTrack. Our results show that with better fitting precision on the clusters, PRIS results yield more robust trajectories after passing through the identical linking step provided in uTrack. Specifically, as compared to the ground truth, uTrack tends to fit the clusters as one particle. The trajectories exhibit little dispersion due to the averaging effect on the locations over all the particles within the cluster. Such bias could influence the follow-up analysis on the diffusion behavior and domain size estimation. On the other hand, spt-PRIS yields separated single particles in the clusters (see zoomed panels in Figure 6) and successfully yields separate trajectories for different particles within the cluster with less dispersion than uTrack results.

**Fig. 6.**
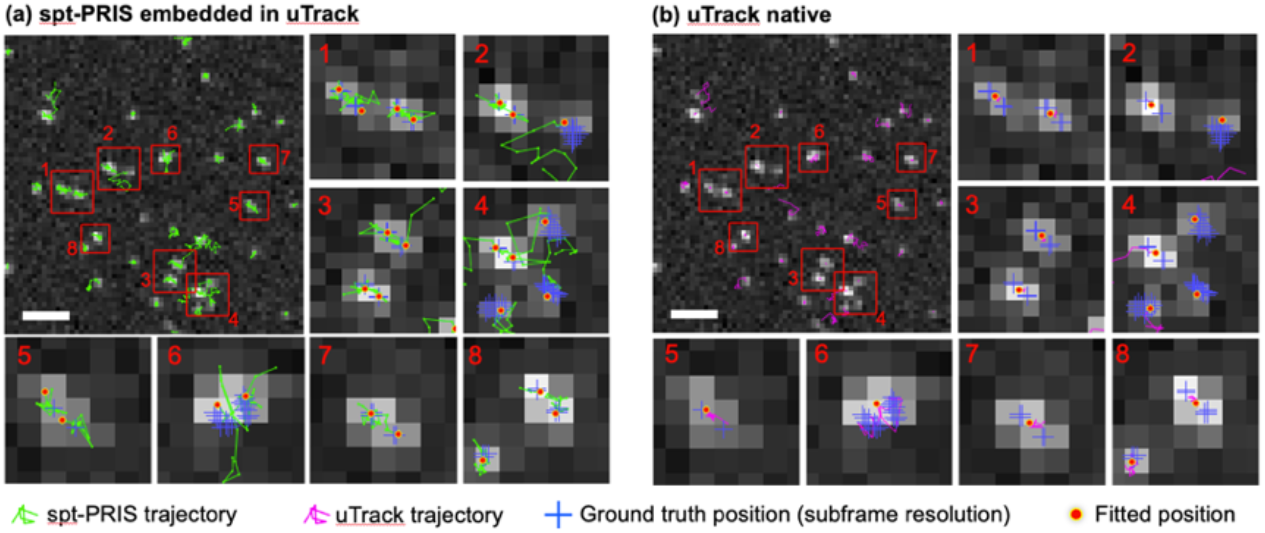
Diffusion trajectories characterization (simulation set 4). (a) shows the result for spt-PRIS, and (b) shows the result for uTrack. Diffusion trajectories for 20 frames are shown, with the particle positions in the last frame annotated as the fitted positions. The ground truth positions contain the motion effect of the 10 subframe positions in the last frame. Scale bars: 16 μm. Pixel size: 160 nm. The complete set of results are shown in visualization 3.

### 3.3 spt-PRIS yields more robust analysis on diffusion

For SPT, further analysis is required on the tracking trajectories to study the diffusion behavior of the molecule of interest. In particular, we validate that the improved fitting and tracking in spt-PRIS is sufficient to improve the robustness of downstream analysis on diffusion states when the particles diffuse in membrane domains. Here, we simulate particles diffusing in membrane domains with a diameter of 200 nm and compare the results of spt-PRIS and uTrack by comparing the global mean square displacement (MSD) as a function of delay times (MSD-Δt plots). Each simulation contains 25 compartments with a radius of 200 nm and a given number of particles randomly spawned into the compartments with specified diffusion constants with confined diffusion.

We first examine the performance with various diffusion constant (0.3 μm^2^/s, 0.1 μm^2^/s, 0.07 μm^2^/s, and 0.03 μm^2^/s) relevant to the reported KRAS4b diffusion states with confinement (Simulation Set 5) [6]. Figure 7 shows the MSD-Δt curves for spt-PRIS, uTrack, and the ground truth. For the medium and fast diffusion speeds (D = 0.07 μm^2^/s, 0.1 μm^2^/s, and 0.3 μm^2^/s), spt-PRIS results are closer to the ground truth than the uTrack results. In the case of D = 0.03 um^2^/s, the spt-PRIS result exhibit more bias than uTrack. Note here that there is bias in the form of more-prominent confinement in all cases with larger Δt, suggesting that error cancelation could exist in the case with D = 0.03 μm^2^/s in the uTrack results where the diffusion constant were initially biased to be faster than the ground truth with lower Δt values, and the universal reduction of MSD values at higher Δt values could cancel the initial bias and drive the MSD closer to the ground truth.

**Fig. 7.**
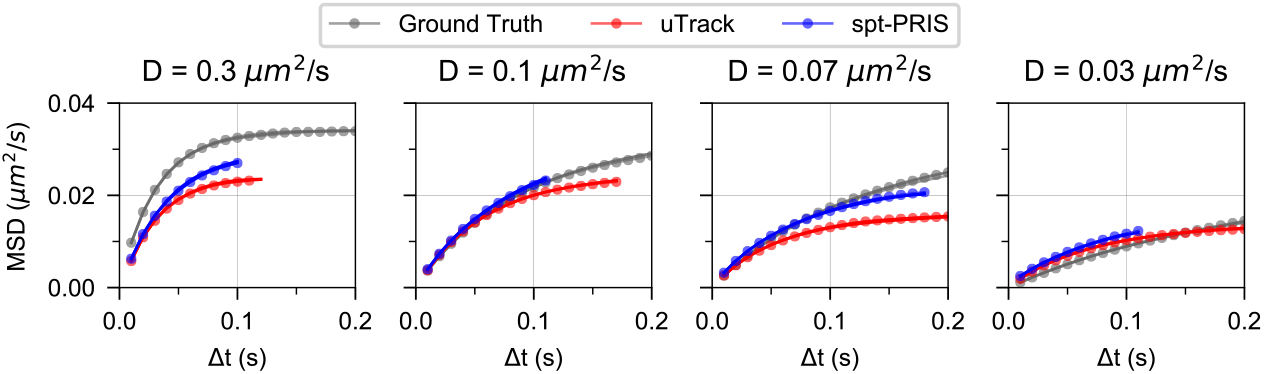
Simulation validation on confined diffusion with different diffusion speeds (simulation set 5). The spt-PRIS results are closer to the ground truth in the medium diffusion speeds, as shown in (b) and (c). Both methods demonstrate biased confinement in the fast diffusion speed shown in (a) and (d).

We then study the influence of cluster size on the extracted diffusion statistics (simulation set 6). In this case, the instantaneous counts of particles (N) were maintained constant (with N = 20, 40, 60, or 80) but still randomly spawned into 25 compartments with a radius of 200 nm. Additional particles were spawned at random when particles dissociate from the membrane (end of the trajectories) to maintain a constant N. This way, the number of particles within each compartment still fluctuates but maintains the nonuniform and uncontrolled distribution of clustering dynamics expected in actual experiments. We can see from Figure 8 that the performance of spt-PRIS is better than that of uTrack in all simulated cases.

**Fig. 8.**
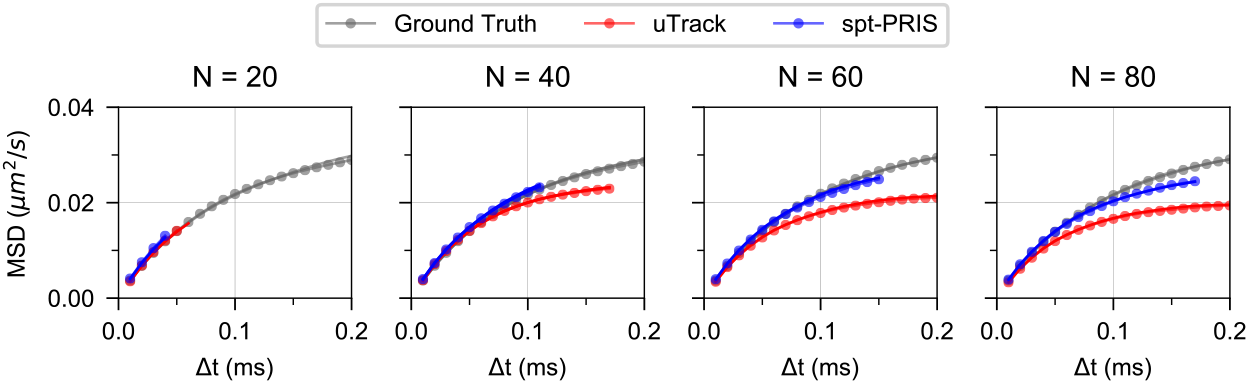
Simulation validation on confined diffusion with different cluster size (simulation set 6). MSD-dT plots are shown for spt-PRIS, uTrack, and the ground truth. In all cases simulated in this set of simulations, the spt-PRIS results are closer to the ground truth than the uTrack results.

We next examine a more complicated case with particle splitting and merging dynamics in the last set of simulations (simulation set 7). As shown in Figure 9(a), each pair of particles is initiated as two monomers that exhibit free diffusion with D=1 μm^2^/s. The monomer particles can enter the R=200 nm compartments of membrane domains and experience a reduced diffusion constant of 0.5 μm^2^/s, from which the pair of particles could escape and restore the faster diffusion constant of 1 μm^2^/s, or merge into dimers where both will share a diffusion trajectory with even slower diffusion constant (0.01 μm^2^/s), after which the dimers will dissociate from the membrane (simulating the loss of fluorescence signal). We compare the MSD-Δt curves for spt-PRIS results, uTrack results, and ground truth. Figure 9(b) shows that MSD-Δt curves from spt-PRIS analysis are closer to the ground truth than that from the uTrack analysis, suggesting spt-PRIS is more robust than uTrack.

**Fig. 9.**
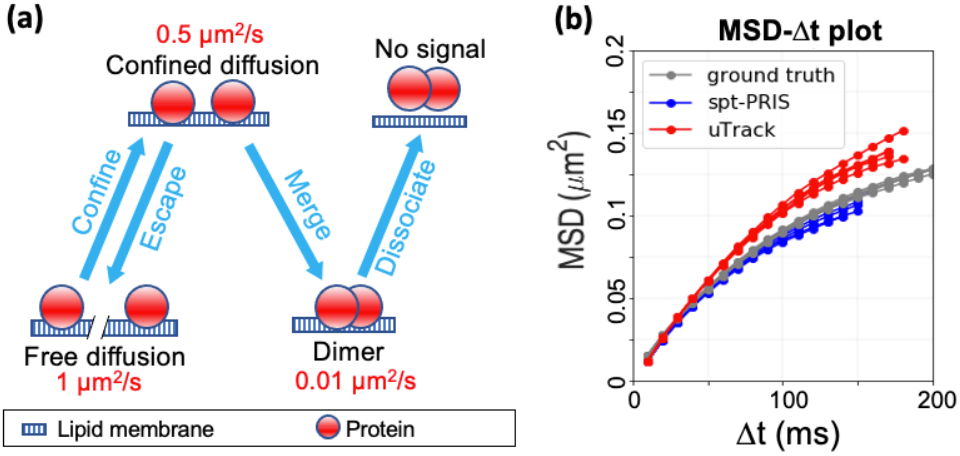
Complex clustering dynamics simulation (simulation set 7). (a) shows the four-state transition of diffusing monomers with either free diffusion or confined diffusion, followed by dimerization and dissociation. (b) shows the MSD-Δt plot from spt-PRIS, uTrack, and the ground truth. We note that spt-PRIS results are closer to the ground truth in all five repetitions in the simulations.

## 4. Application and comparison on *in-vitro* KRAS4b dynamics measurements

Finally, we apply spt-PRIS to the study of KRAS4b protein dynamics in comparison with uTrack on supported lipid bilayers with two different lipid compositions: First, a simple 2-lipid SLB of zwitterionic POPC (87%) and negatively charged POPS (13%). Second, a more complex 8-lipid SLB composed of eight different lipid classes representing the most abundant lipid species found in the inner leaflet of the plasma membrane of mammalian cells [19,27]. More pronounced KRAS4b clustering effects are expected on the 8-lipid SLB than the simple 2-lipid SLB due to negatively charged lipids [19,27]. 4 movies for the sample with an 8-lipid SLB, and 13 movies for the sample with a 2-lipid SLB were analyzed. Tracking graph (TG) analysis was used to remove the linking imperfections in the trajectories and to classify features into subpopulations based on different TG size bounds (*S*_*b*_), which estimates a lower bound on the number of particles of a particular feature [17]. Note here that the *S*_*b*_ is related but does not directly represent the number of labeled KRAS4b proteins. Instead, the total number of KRAS4b molecules for a cluster could be higher than the *S*_*b*_ because of the unlabeled fraction or the dark population, while the total number of bright KRAS4b molecules could be lower than the *S*_*b*_ due to undetected dissociation or bleaching events. The statistical analysis was performed on each subpopulation over all the analyzed movies.

As shown in Figure 10(a)(b), in both 2-lipid and 8-lipid SLBs, spt-PRIS yields significantly more trajectories than the uTrack results, providing more statistics for the subsequent analyses, especially on the subpopulations of clusters with TG size bounds greater than 1. uTrack provided substantial trajectory counts for *S*_*b*_=1 and *S*_*b*_=2 for the 8-lipid SLB (Figure 10(b)) and very sparse counts for *S*_*b*_=2 on the 2-lipid SLB (Figure 10(a)). The importance of the trajectory abundance is clearly demonstrated in the robustness of the MSD-Δt curves calculated for each subpopulation using both spt-PRIS and uTrack analysis, as shown in Figure 10(c).

**Fig. 10.**
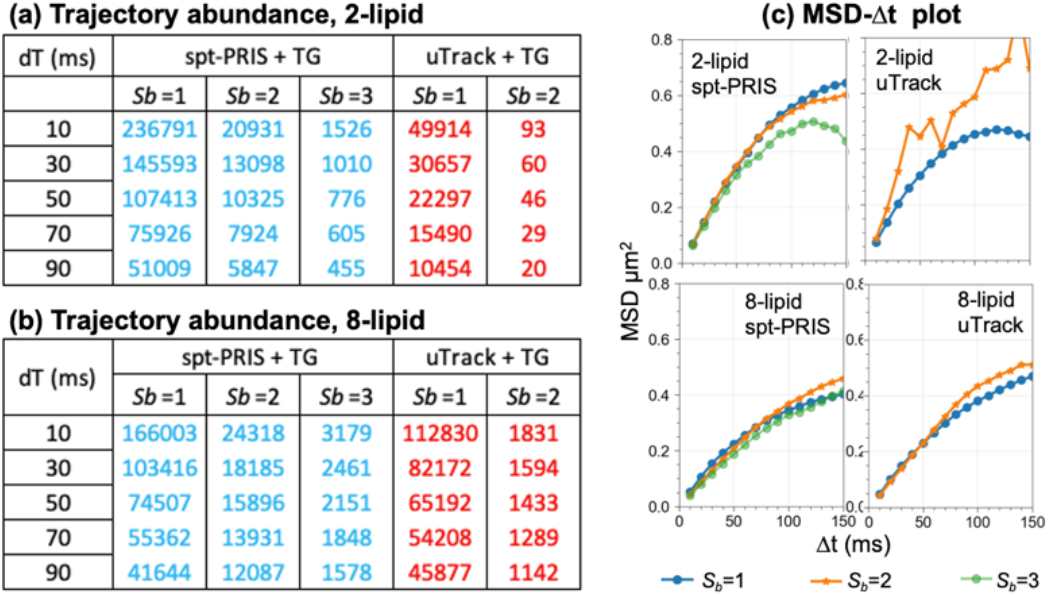
MSD trajectory abundance and MSD-Δt plots for experimental data. (a) and (b) list out the number of trajectories for each condition as labeled in the tables. (c) shows the MSD-Δt plot for the annotated conditions.

We further compared the performance of spt-PRIS to the uTrack by subjecting the trajectories to three types of statistical analysis: mean square displacement (MSD) distribution (see section 4.1), brightness distribution (see section 4.2), and Jumping Distance (JD) analysis for diffusion states (see section 4.3) [30–32]. In section 4.4, we also discuss the unexpected yet interesting result for KRAS4b on the 2-lipid SLB where two distinct subpopulations are identified from the spt-PRIS results followed by TG analysis while the diffusion states are the same.

### 4.1 MSD distribution

MSD distributions of the spt-PRIS and uTrack results are shown in Figure 11 for the *S*_*b*_=1 and *S*_*b*_=2 subpopulations for both lipid compositions. Here, the MSD values were calculated for each trajectory, and the distributions were calculated over all datasets. Both methods demonstrate robust distributions for the *S*_*b*_=1 subpopulation; however, in the case of *S*_*b*_=2, spt-PRIS results showed significantly more robust distribution.

**Fig. 11.**
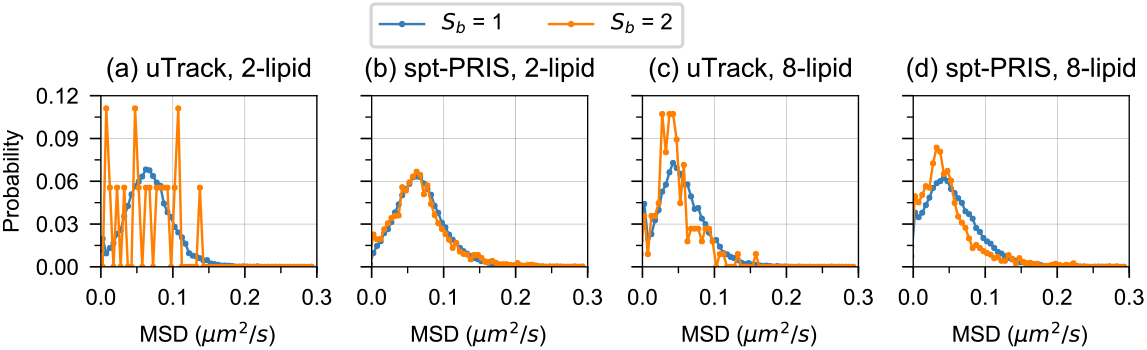
MSD distributions for experimental data with TG size bounds of 1 and 2. The results for 2-lipid and 8-lipid SLB data from both uTrack and spt-PRIS analysis are shown as annotated. The probability distribution of MSD is normalized to have total probability of 1. The MSD values are calculated with time delay of 1 movie frame (Δt =10 ms).

For the 2-lipid SLB (Figure 11(a)(b)), the uTrack results demonstrate a dominant population of KRAS4b clusters with *S*_*b*_=1, whereas the spt-PRIS results show two subpopulations for *S*_*b*_=1 and *S*_*b*_=2, which share nearly identical distributions. The weak distinction between the two MSD distributions indicates homogeneous diffusion of KRAS4b on the 2-lipid SLB independent of the size bound. For the 8-lipid SLB (Figure 11(c)(d)), both methods yield significant subpopulations with *S*_*b*_=1 and *S*_*b*_=2. However, due to the more significant number of trajectories obtained from spt-PRIS, the MSD distribution exhibits reduced variation compared to uTrack. For both methods, the MSD distribution profiles for *S*_*b*_=2 are skewed towards the smaller MSD values when compared to the *S*_*b*_=1 case, suggesting the two subpopulations have different diffusion behavior. This observation agrees well with the prior studies where KRAS4b proteins exhibited more prominent clustering dynamics in 8-lipid SLB.

### 4.2 Brightness distribution

Figure 12 shows the analysis of brightness distributions for the *S*_*b*_=1 and *S*_*b*_=2 subpopulations for both lipid compositions. For the 2-lipid SLB, uTrack identifies one dominant population with *S*_*b*_=1, whereas spt-PRIS identifies *S*_*b*_=1 and *S*_*b*_=2 as two distinct subpopulations. A small difference of the brightness distribution is presented. In the case of 8-lipid SLB data, which has a more pronounced KRAS4b clustering effect based on prior studies [6], both methods demonstrate separation of 2 subpopulations while spt-PRIS demonstrates a more smooth distribution than uTrack. Both exhibit an up-shift of the brightness distribution in the *S*_*b*_=2 subpopulation when compared to the *S*_*b*_=1 subpopulation, confirming the more prominent clustering dynamics in *S*_*b*_=2 population.

**Fig. 12.**
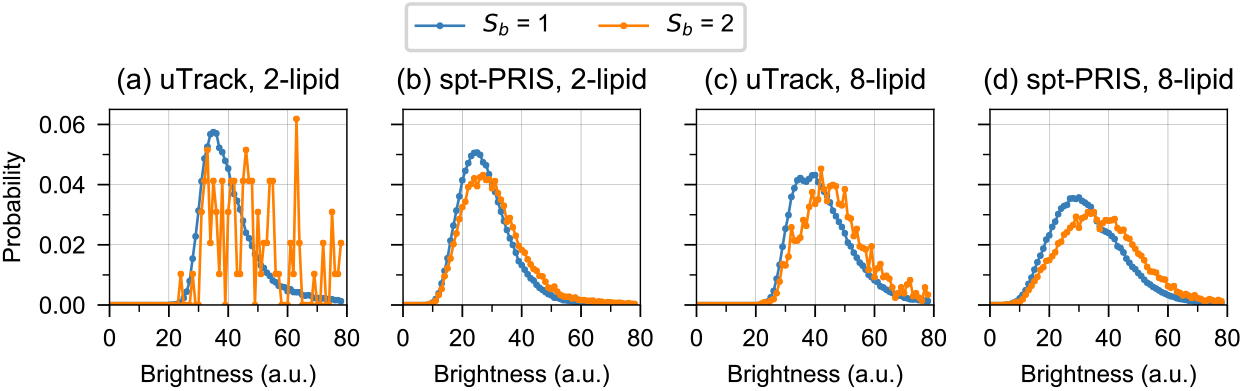
Brightness distribution of features in spt-PRIS results and uTrack results. The results for 2-lipid and 8-lipid SLB data from both spt-PRIS and uTrack analysis are shown as annotated. Arbitrary units were used for brightness.

### 4.3 Jump Distance (JD) analysis

We further analyze the diffusion states of the subpopulations with different TG size bounds using jump distance analysis [30–32]. Least-square fitting was used to fit the cumulative probability density function of the square of the jumping distances (*r*^*2*^) to a mixed species model shown in Equation (1),

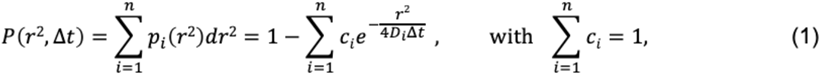

where *r* is the jump distance of the given time interval (Δt), *n* is the total number of species accounted in the fitting model, *i* is the species index, *c*_*i*_ is the fraction coefficient for the *i*^th^ species, *p* is the probability density of the square value of jump distances extracted from the global histogram of the jumping distances over all analyzed movies, and *D* is the diffusion constant. We performed the analysis for both uTrack and spt-PRIS results on all the subpopulations with *S*_*b*_ values of 1, 2, and 3 on both 2-lipid and 8-lipid SLBs. For each dataset, we performed fitting with *n*-species model shown in Equation (1) with *n* ranging from 1 to 6, and the best fit is determined by the highest R^2^ value characterizing the fitting quality. To avoid over-fitting, the results with species fractions lower than 1% or with species sharing identical diffusion constants are discarded.

The best fit results are shown in Figure 13. For 2-lipid SLB data, the spt-PRIS results for all the subpopulations and the uTrack results for *S*_*b*_=1 subpopulation identifies similar diffusion constants (1.83 to 1.88 μm^2^/s) with a 1-species model, consistent with the MSD distribution analyses shown in Section 4.1 and further confirming a single composition of diffusion state for KRAS4b on the 2-lipid SLB. However, uTrack results for *S*_*b*_=2 identifies a single species with a diffusion constant of 2.57 μm^2^/s, which we assess as unreliable due to the lower R^2^ value, and the much larger range of fitting residuals when compared to the other fitting results. The uTrack result for the *S*_*b*_=3 subpopulation is insufficient to support the JD analysis. For the 8-lipid SLB data, the spt-PRIS results for all the subpopulations fit best to a 3-species model with slow, medium, and fast diffusion species. The diffusion coefficients for individual states remained similar across all the subpopulations with *S*_*b*_ ranges from 1 to 3. With increasing *S*_*b*_, the composition of the subpopulation shifts toward the slower diffusion state, with either a slower diffusion constant for the slow and medium diffusion species (when *S*_*b*_ increases from 1 to 2) or decreasing fractions of the fast-diffusing species and increasing fractions of the slower-diffusion species (when *S*_*b*_ increases from 2 to 3). However, the uTrack analysis results best fit a 2-species model for the *S*_*b*_=1 subpopulation, and best fits a 1-species model for the *S*_*b*_=2 and *S*_*b*_=3 subpopulations, which is inconsistent with the 3-species model for 8-lipid SLB in the prior studies [6].

**Fig. 13.**
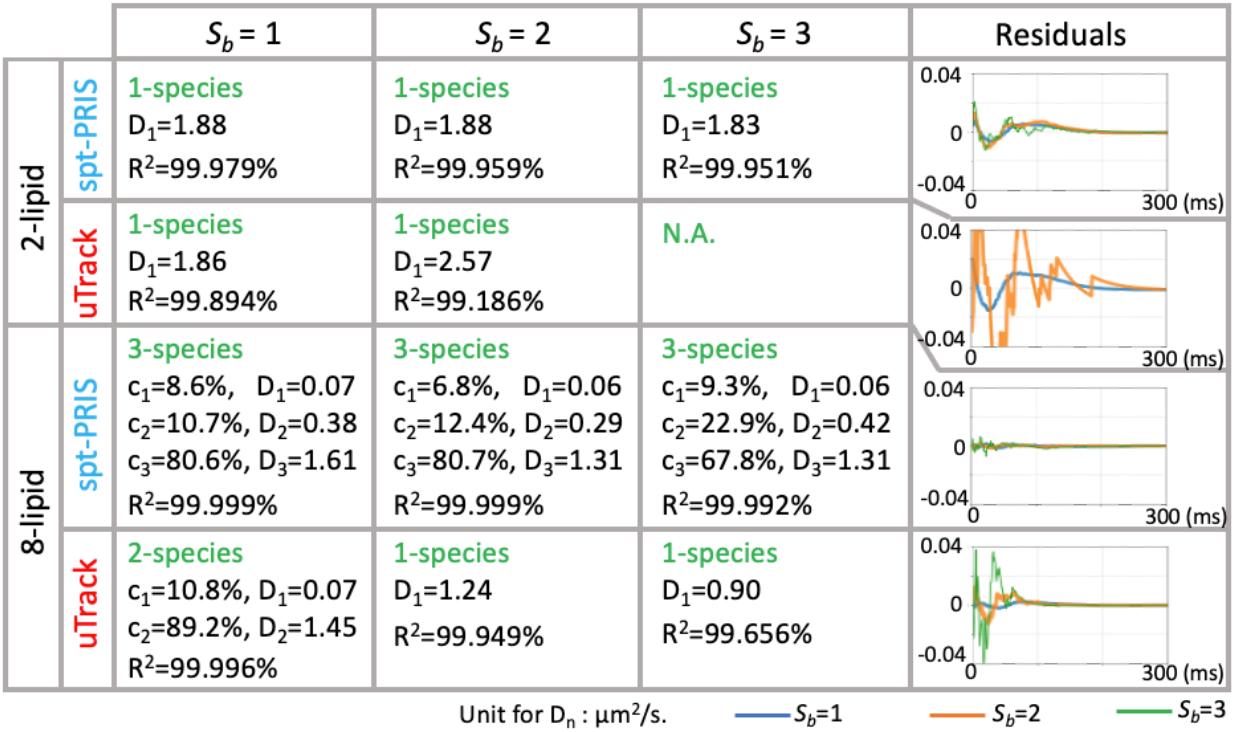
Diffusion state analysis on 8-lipid data. We attempted the fitting to Equation (1) with n-species models (n ranges 1 to 6) for all the data. The best-fit results are shown with the n^th^ species characterized by its fraction c_i_ and diffusion constant D_i_, where i is the species index. The R^2^ score of each fitting is displayed for each condition, and the fitting residuals for each method are shown in the last column. The unit for the diffusion constants is μm^2^/s.

For all the cases, the spt-PRIS results yield more robust JD analysis fittings than the uTrack results, with lower amplitude of fitting residuals and higher R^2^ score. By providing access to the robust statistical analyses into the subpopulations with higher *S*_*b*_ values, spt-PRIS provides more information in understanding the KRAS4b dynamic properties.

### 4.4 Distinct KRAS4b subpopulations in the 2-lipid supported lipid bilayer

While we expect 2-lipid data to demonstrate less complex diffusion behavior as compared to the 8-lipid data due to the lack of countercharged lipids, the spt-PRIS results identify two distinct subpopulations with *S*_*b*_=1 and *S*_*b*_=2 respectively in the 2-lipid data, in contrast with the uTrack results, which identify a single dominant *S*_*b*_=1 population from the dataset. No differences in diffusion states were found from the MSD distribution (Figure 11(b)) and the jump distance analysis (Figure 13). However, distinctions in the brightness distribution are shown in Figure 12(b), similar to that from the 8-lipid data (which could serve as a positive control) from both spt-PRIS and uTrack results shown in Figure 12(c)(d).

Here we discuss the unexpected *S*_*b*_=2 subpopulation from spt-PRIS (combined with TG analysis) results on the 2-lipid data. First, sufficient abundance of the trajectories is shown with larger Δt values (Figure 10(a)), suggesting that it is not representing the crossing-over of independently diffusing bright particles where the PSFs would only overlap transiently. Second, the MSD-Δt plots shown in Figure 10(c) demonstrate significant long-range displacements with larger Δt values, suggesting that it is less likely to be due to the nonspecific binding of labeled KRAS4b on the coverslip, which would appear more immobile with smaller MSD at greater Δt. Third, slightly more prominent confinement of this population is shown when compared to the *S*_*b*_=1 subpopulation (Figure 10(c), 2-lipid spt-PRIS), suggesting a weak clustering effect. We hypothesize that such an *S*_*b*_=2 subpopulation in the 2-lipid SLB houses a distinct state. This state could be seen as ‘attempts’ of KRAS4b to cluster but without being able to form clusters due to the lack of countercharge lipids required for stabilization. The distances between the particles could have dropped below the resolution limit, which contributes to the increase of brightness when multiple particles are fitted as one and is only distinguishable with the higher resolution limit provided by PRIS than the conventional method. Nonetheless, we acknowledge that more in-depth theoretical analyses, modeling studies, and in-vitro experiments are required to confirm such a hypothesis and to gain additional biological insights.

## 5. Discussion

In this work, we developed “spt-PRIS” --an enhanced SPT method with improved localization using compressive sensing. Our method adopts PRIS to solve the compressive sensing problem and integrates it into a state-of-the-art open-source tool, uTrack, to leverage its linking algorithm and enhance its ability to capture merging and splitting events. We performed systematic simulations to characterize and validate the improved SPT performance using spt-PRIS, especially for datasets where resolving overlapping PSFs is critical. We applied spt-PRIS to experimental data with KRAS4b clustering dynamics on 2-lipid and 8-lipid membrane SLB and analyzed the tracking graph for trajectory cleaning and subpopulation classification prior to diffusion state analysis on each subpopulation. Our results demonstrate that spt-PRIS provides an unprecedented SPT capability to study particle interactions where close-range interactions are critical. spt-PRIS also provides previously inaccessible statistics for the subpopulation analysis when combined with tracking graph analysis, bringing in extra information into the study of the KRAS4b system.

Further development of spt-PRIS includes adopting advanced algorithm architecture or computation platforms to improve efficiency, such as implementation in Halide, a domain-specific language for image processing [33], implementation on field programmable gate array (FPGA), graphic processing unit (GPU), or integrating with cloud computing. The spt-PRIS method also requires improvement of the method accessibility through developing a user-friendly and open-source toolbox. Integration of spt-PRIS with state-of-the-art imaging modalities is also plausible, for example, with high-resolution light sheet microscopes such as lattice light sheet microscope (LLSM) [23] and other light sheet imaging systems [34,35], and point spread function engineering [36–39] to further engage the multi-channel, multi-species and multi-dimensional localization potential of PRIS reconstruction. Additionally, our work motivates further development of linking and classification algorithms for SPT with awareness of motion effect and features with dynamic close-range interactions. Our work also motivates using spt-PRIS in different subjects of study as an SPT tool to reach a broader range of interests in the scientific community.

## Supporting information

visualization 1

visualization 2

visualization 3

## Acknowledgements

This work has been supported in part by the Joint Design of Advanced Computing Solutions for Cancer (JDACS4C) program established by the U.S. Department of Energy (DOE) and the National Cancer Institute (NCI) of the National Institutes of Health (NIH). This work was performed under the auspices of the U.S. DOE by Lawrence Livermore National Laboratory under Contract DE-AC52-07NA27344 and The Frederick National Laboratory for Cancer Research under Contract HHSN261200800001E. Release number: LLNL-JRNL-819405.

